# Distinct mechanisms are employed by T-cell-instructed myeloid cells for IL-1β production in humans and mice

**DOI:** 10.1101/2025.02.26.640341

**Authors:** Jinyi Zhao, Zhu Liang, Benedikt M Kessler, Liye Chen

**Affiliations:** Botnar Research Centre, Nuffield Department of Orthopaedics, Rheumatology and Musculoskeletal Sciences, University of Oxford, Oxford, UK; Chinese Academy for Medical Sciences Oxford Institute, Nuffield Department of Medicine, University of Oxford, Roosevelt Drive, Oxford OX3 7FZ, UK; Target Discovery Institute, Centre for Medicines Discovery, Nuffield Department of Medicine, University of Oxford, Roosevelt Drive, Oxford OX3 7FZ, UK; Department of Microbiology, Boston, MA, USA Division of Infectious Diseases, Brigham and Women’s Hospital, Boston, MA, USA

## Abstract

Interleukin-1 beta (IL-1β) is known as an inflammasome-dependent pro-inflammatory cytokine that has been implicated in T-cell-driven autoimmune diseases. In mice, T-cells were reported to instruct myeloid cells to produce IL-1β via an inflammasome-independent mechanism, that engages TNFR and Fas-caspase-8-dependent signalling pathways. In this study, we explored T-cell-driven myeloid IL-1β production in humans. Co-culturing of autologous primary T-cells (memory CD4+ and CD8+) with myeloid cells (monocyte, macrophage and dendritic cells) revealed that both memory CD4+ and CD8+ T cells induce IL-1β secretion. Also, caspase-1 rather than caspase-8 cleaves pro-IL-1β in human myeloid cells. This process depends on TNF-α, CD40L and IFN-γ together rather than TNF-α alone, leading to upregulated pro-IL-1β expression in human myeloid cells. We additionally show that TNF-α, CD40L and IFN-γ independently enhance IL-1β secretion. Together, our study highlights that, despite the shared biology in T-cell-instructed IL-1β production between humans and mice, different underlying molecular pathways are implicated.

**Summary:** Like murine cells, human T-cells stimulate IL-1β secretion from myeloid cells. However, the underlying mechanisms differ, with IL-1β production in humans being driven by TNF-α, CD40L, IFN-γ, and Caspase-1, whereas in mice, it is primarily regulated by TNF-α and Caspase-8.

## INTRODUCTION

Interleukin-1 beta (IL-1β) is a critical pro-inflammatory cytokine that plays a key role in various physiological and pathological processes (Gabay et al., 2010). It enhances immune responses by promoting inflammation in response to infections and tissue injury (Mantovani et al., 2019). IL-1β is produced predominantly by activated myeloid cells, including blood monocytes, macrophages, and dendritic cells (DCs) (Dinarello et al., 2012). The synthesis and secretion of IL-1β occur through a two-step process. First, pro-IL-1β is synthesized as an inactive precursor. It is then cleaved by enzymes such as caspase-1 or caspase-8 to generate the biologically active cleaved IL-1β (Broz and Dixit, 2016, Agostini et al., 2004, Afonina et al., 2015). Once secreted, IL-1β binds to the IL-1 receptor (IL-1R) on the surface of target cells, initiating downstream signalling pathways that mediate inflammation, fever, and tissue repair (Kolb et al., 2001, Mantovani et al., 2019). Dysregulation of IL-1β is implicated in the pathogenesis of various chronic inflammatory and autoimmune diseases, including rheumatoid arthritis (RA) and inflammatory bowel disease, making it a promising target for therapeutic intervention (Coccia et al., 2012, Dinarello, 2011). IL-1β also modulates immune cell functions by enhancing T cell activation and survival and promoting the effector responses of dendritic cells, macrophages, and neutrophils (Coccia et al., 2012).

The interaction between myeloid cells and T cells is critical for initiating and regulating adaptive immune responses. Myeloid cells activate T cells by presenting processed antigens to T-cell receptors (TCRs) via major histocompatibility complex (MHC) molecules on their surface (Rossjohn et al., 2015). This interaction extends beyond antigen presentation and is further modulated by co-stimulatory signals and cytokines. For instance, the binding of CD40 on myeloid cells to CD40 ligand (CD40L) on T cells enhances both antigen presentation and cytokine production, thereby promoting T-cell activation and differentiation (Schoenberger et al., 1998, Grewal and Flavell, 1996, Karnell et al., 2019). Additionally, cytokines such as TNF-α and IFN-γ, secreted by myeloid cells and T cells respectively, amplify this interaction and drive inflammatory responses. These cytokines upregulate co-stimulatory molecules such as CD80 and CD86, as well as adhesion molecules on myeloid cells, strengthening their interactions with T cells (Psarras et al., 2021, Chen et al., 1998, Schroder et al., 2004).

IL-1β has been implicated in T cell-mediated autoimmunity, including rheumatoid arthritis (RA) and juvenile idiopathic arthritis (JIA), prompting the development of IL-1β-blocking antibodies for these conditions (Schett et al., 2016). The precise mechanism underlying IL-1β production in T cell-mediated autoimmunity was unclear until a recent study demonstrated inflammasome-independent IL-1β production in myeloid cells stimulated by effector CD4+ T cells using murine models (Jain et al., 2020). Specifically, this study revealed that pro-IL-1β synthesis was driven by TNF-α signalling, while its cleavage into the active form relied on the Fas/Caspase-8 pathway. In our study, we investigated whether a similar mechanism operates in humans. Our findings confirm that activated T cells can induce IL-1β secretion by human myeloid cells, but the underlying mechanisms differ significantly from those observed in murine cells. Unlike the exclusive reliance on TNF-α signalling in mice, pro-IL-1β expression in human cells is governed by three non-redundant signals from activated T cells: TNF-α, CD40L, and IFN-γ. Furthermore, we show that pro-IL-1β cleavage in human cells depends on Caspase-1, rather than the Fas/Caspase-8 pathway. Together, our findings, along with the previous murine study, underscore species-specific differences in the molecular pathways regulating IL-1β production in T cell-mediated autoimmunity. These insights provide a better understanding of IL-1β regulation and its role in autoimmune diseases.

## RESULTS AND DISCUSSION

### Activated T cells drive IL-1β production by human primary monocytes

Monocyte infiltration is a hallmark of inflammation in autoimmune diseases. These recruited monocytes constitute a significant portion of the infiltrating myeloid cells and interact with activated T cells in the inflammatory milieu. We observed IL-1β secretion by monocytes upon stimulation with activated autologous memory CD4+ T cells (mCD4) (Fig. 1A). Importantly, T cell activation was crucial for this response and induced IL-1β production in a dose-dependent manner (Fig. 1B).

**Figure 1.**
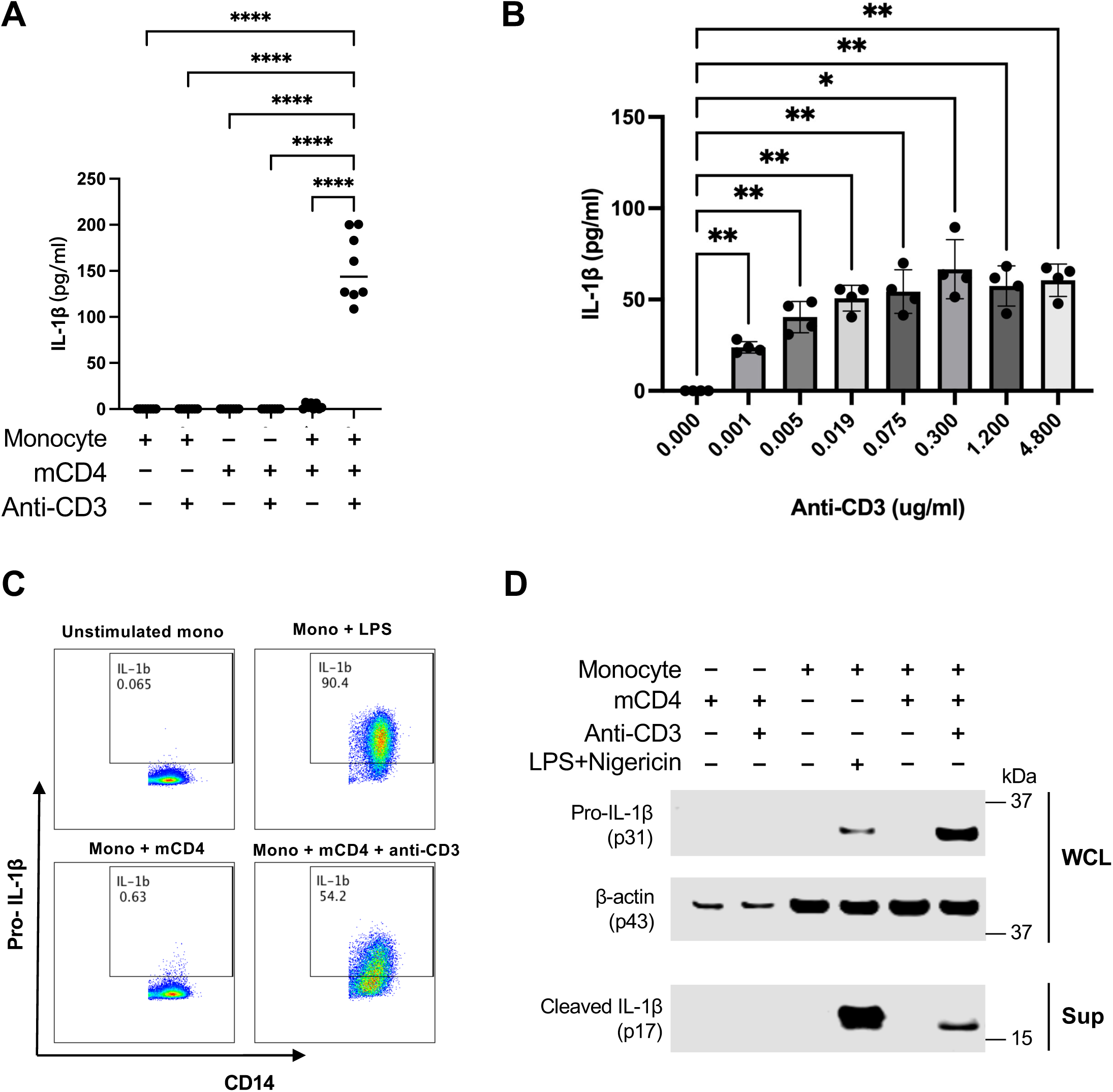
T-cell activation triggers the production and secretion of IL-1β in monocytes. (A) Isolated monocytes and autologous memory CD4+ T-cells from human PBMCs (n=8) were cultured alone or co-cultured in the absence/presence of anti-CD3 antibodies for 16 hours. The levels of cytokine IL-1β in the culture supernatant were measured using ELISA. (B) Isolated monocytes from human PBMCs (n=4) were co-cultured with autologous memory CD4+ in the presence of graded concentrations of anti-CD3 antibodies for 16 hours. The levels of cytokine IL-1β in the culture supernatant were measured using ELISA. (C) Representative flow cytometry plots and summary graphs showing the expression of intracellular pro-IL-1β by isolated CD14+ monocytes from human PBMCs simulated with LPS or co-cultured with autologous memory CD4+ T-cells in the absence/presence of anti-CD3 antibodies for 16 hours. (D) Immunoblotting analysis of pro-IL-1β, β-actin, and cleaved-IL-1β in the whole cell lysates (WCL) or the supernatants from co-culture systems. Data are represented as mean and SEM of independent donors in independent experiments. The P-value was assessed by one-way ANOVA (* P≤ 0.05; ** P≤ 0.01; *** P≤ 0.001).

We then explored the effects of T cell activation on pro-IL-1β and cleaved IL-1β expression in monocytes. Intracellular staining of monocytes revealed a robust induction of pro-IL-1β expression upon the addition of an anti-CD3 antibody to the monocyte and mCD4 co-culture (Fig. 1C and Fig. S1A). Consistent with these findings, immunoblotting of whole cell lysates (WCL) from the monocyte and mCD4 co-culture with anti-CD3 stimulation confirmed the presence of pro-IL-1β (Fig. 1D). Furthermore, IL-1β activation was validated by detecting cleaved IL-1β in the supernatant (Sup) (Fig. 1D). Since CD8+ T cells also play a significant role in autoimmune responses, we investigated whether activated memory CD8+ T cells (mCD8) could similarly induce IL-1β production in monocytes. We observed comparable results (Fig. S2). Collectively, these data demonstrate that activated T cells, both CD4+ and CD8+, can induce IL-1β secretion by human primary monocytes.

### IL-1β cleavage in T-cell-instructed monocyte activation depends on caspase-1 rather than caspase-8

Caspase-1 is the primary enzyme responsible for cleaving pro-IL-1β into its active form, IL-1β. Interestingly, prior studies demonstrated that Caspase-8, rather than Caspase-1, mediated pro-IL-1β activation in murine myeloid cells stimulated by T cells (Jain et al., 2020). In our study, we observed that siRNA-mediated knockdown of Caspase-1, but not Caspase-8, in human monocytes significantly reduced IL-1β secretion induced by T-cell activation (Fig. 2A). Importantly, this effect is unlikely due to differences in knockdown efficiency, as both Caspase-1 and Caspase-8 protein levels were suppressed in a comparable fashion (Fig. S3). We then used the well-characterized caspase-1 inhibitor (Ac-YVAD-cmk) and caspase-8 inhibitor (Z-IETD-FMK) for additional validation (Corasaniti et al., 2003, Zhang et al., 2016, Thornberry et al., 1997, Komoriya et al., 2000). In line with the findings from siRNA knockdown experiments, a significant reduction of secreted IL-1β induced by T-cell stimulation was observed in monocytes pre-treated with inhibitors for caspase-1 but not caspase-8 (Fig. 2B).

**Figure 2.**
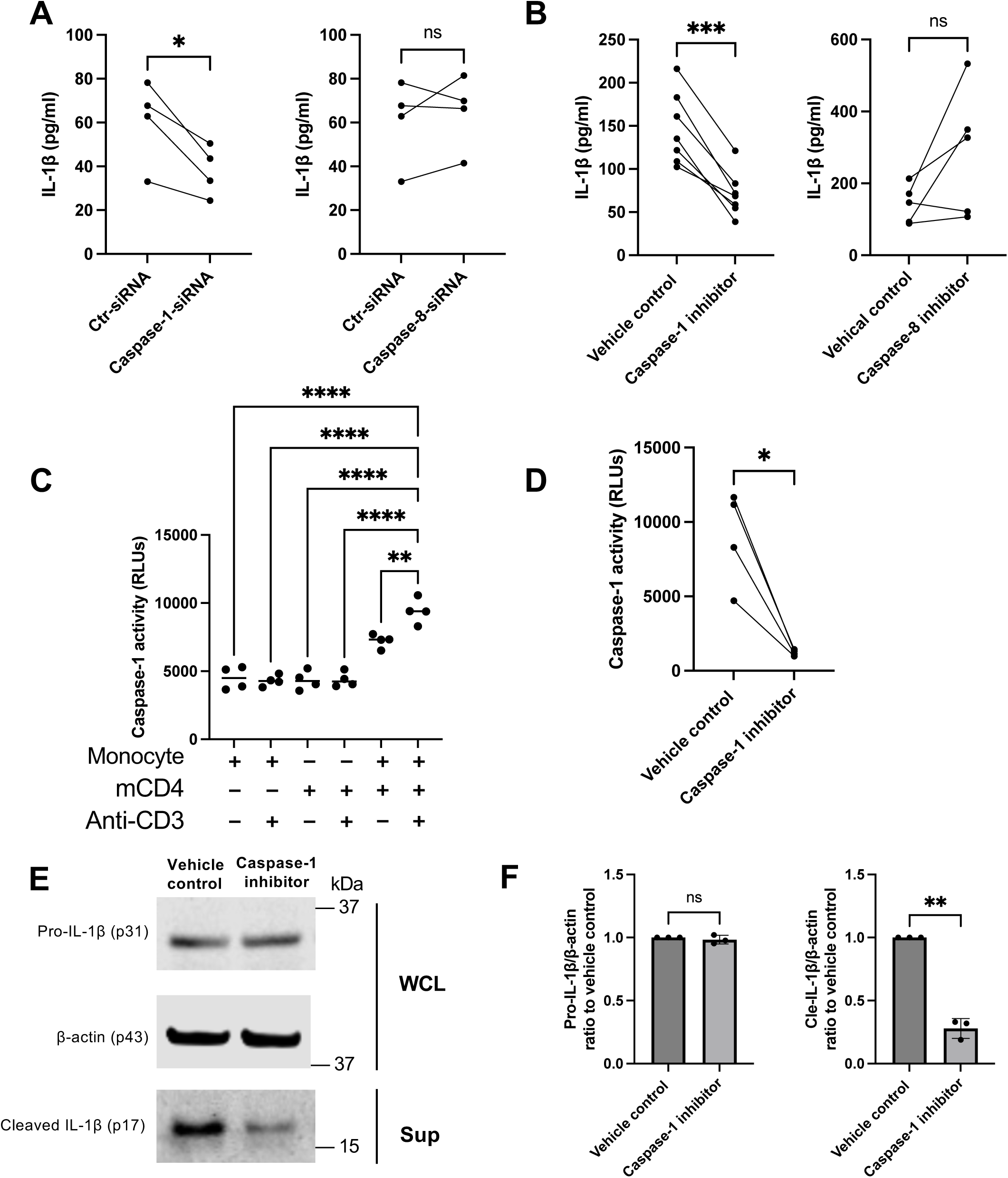
The cleavage of IL-1β during T-cell-induced monocyte activation depends on caspase-1 rather than caspase-8. (A) Isolated monocytes from human PBMCs (n=4) were transfected with non-targeting control siRNA, CASP1 siRNA, or CASP8 siRNA before co-culture with autologous memory CD4+ T cells and anti-CD3 antibodies for 16 hours. The level of IL-1β in culture supernatant was measured by ELISA. (B) Isolated monocytes from human PBMCs were pre-treated with the caspase-1 inhibitor or caspase-8 inhibitor for 4 hours, followed by co-culture with autologous memory CD4+ T-cells and anti-CD3 antibodies for 16 hours. The level of IL-1β in culture supernatant was measured by ELISA. (C) Caspase-1 activity in the supernatant was measured using the Caspase-Glo® 1 Inflammasome Assay (Promega) and expressed as relative luminescent units (RLUs). (D) Isolated monocytes from human PBMCs were pre-treated with the caspase-1 inhibitor for 4 hours, followed by co-culture with autologous memory CD4+ T-cells and anti-CD3 antibodies for 16 hours. Caspase-1 activity in the supernatant was measured using the Caspase-Glo® 1 Inflammasome Assay (left); The level of IL-1β in the culture supernatant was measured by ELISA (right). (E) The protein levels of pro-IL-1β, β-actin, and cleaved-IL-1β in the whole cell lysates or the supernatants from co-culture systems were measured by Western blotting. (F) The ratio of relative protein expression of pro-IL-1β and cleaved-IL-1β (inhibitor-treated vs. vehicle control). Data are represented as mean and SEM of independent donors in independent experiments. The P-value was assessed by students t-test or one-way ANOVA (* P≤ 0.05; ** P≤ 0.01; *** P≤ 0.001).

To further confirm the role of caspase-1 in the IL-1β response, we first measured caspase-1 activity in the supernatant and observed a significant increase following the addition of anti-CD3 to monocyte and mCD4 co-cultures (Fig. 2C). Pre-treatment of monocytes with the caspase-1 inhibitor followed by T cell stimulation significantly reduced caspase-1 activity in the supernatant (Fig. 2D). Immunoblot analysis further revealed that caspase-1 inhibition selectively reduced the levels of cleaved IL-1β without affecting pro-IL-1β, providing strong evidence that caspase-1 is responsible for IL-1β activation in human monocytes stimulated by T cells (Fig. 2E and F).

### TNF-α, CD40L, and IFN-γ, but not Fas ligand, drive IL-1β secretion by promoting pro-IL-1β expression

T-cell-derived TNF-α and Fas ligand (FasL) have been shown to drive the upregulation and cleavage of pro-IL-1β, respectively, in murine myeloid cells stimulated by T-cells (Jain et al., 2020). In addition to TNF-α and FasL, activated T-cells are known to express CD40 ligand (CD40L) and IFN-γ, both of which play key roles in activating myeloid cells within inflamed tissues. To investigate whether these factors influence IL-1β secretion by human primary monocytes stimulated by mCD4 cells, we performed immunoblotting analyses. Blocking TNF-α, CD40L, or IFN-γ, but not FasL, reduced pro-IL-1β expression in whole-cell lysates and decreased cleaved IL-1β levels in the supernatant (Fig. 3A and B). Interestingly, the ratio of cleaved IL-1β to pro-IL-1β remained unaffected by any of the blocking antibodies, suggesting that these factors regulate pro-IL-1β expression rather than its cleavage. Consistent with this, caspase-1 activity in the supernatant was unaltered by the blocking antibodies (Fig. 3D).

**Figure 3.**
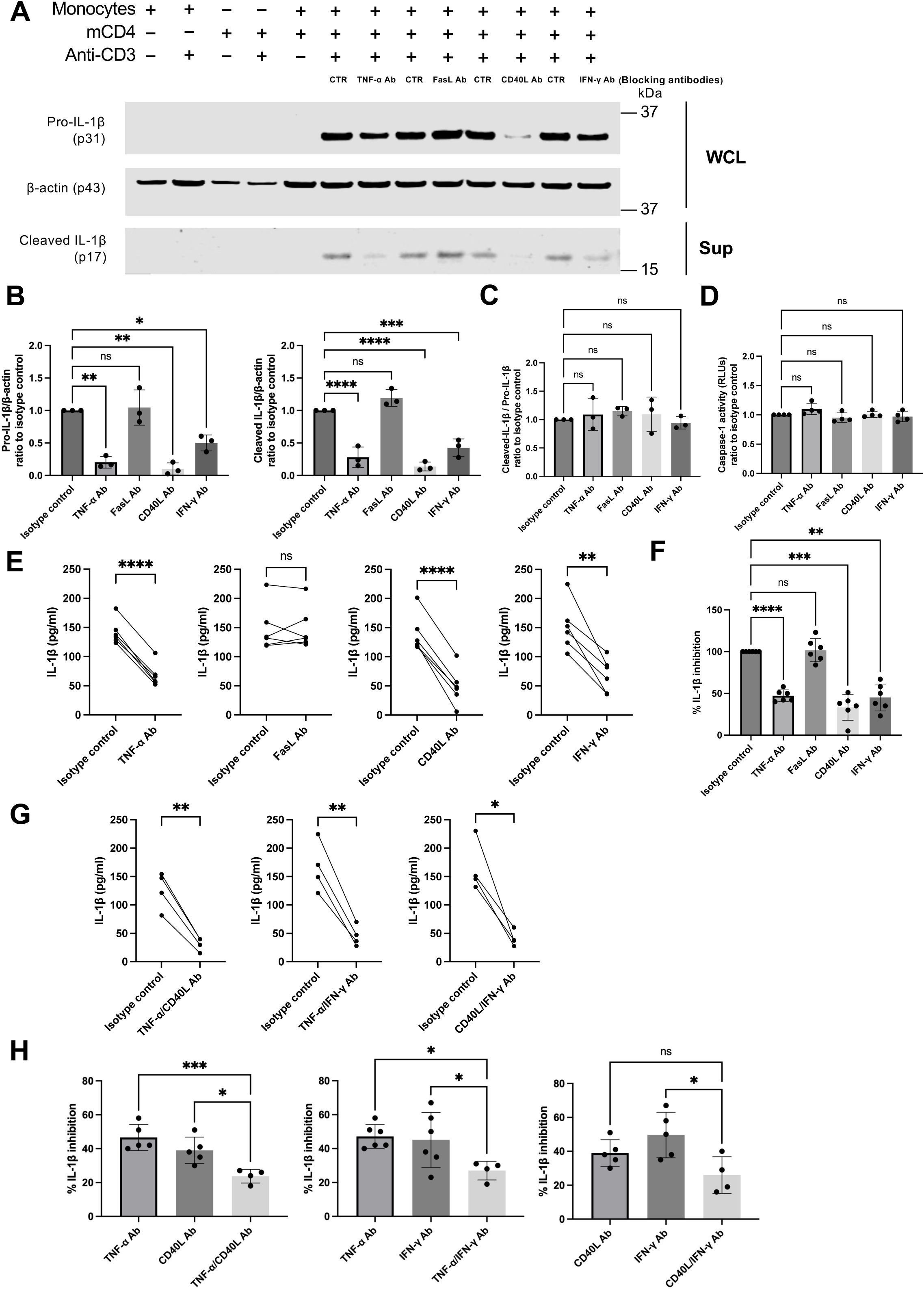
TNF-α, CD40L, and IFN-γ, but not Fas ligand, drive the secretion of IL-1β through the upregulation of pro-IL-1β. Isolated monocytes and autologous memory CD4+ T-cells from human PBMCs were cultured alone or co-cultured in the absence/presence of anti-CD3 antibodies and neutralising antibodies against TNF-α, FasL, CD40L, and IFN-γ for 16 hours. (A) Immunoblotting analysis showed the protein expression of pro-IL-1β, β-actin, and cleaved-IL-1β in the whole cell lysates (WCL) or the supernatants from co-culture systems. (B) The ratio of the relative protein expression of pro-IL-1β and cleaved-IL-1β in neutralising antibody-treated samples to isotype control-treated samples. (C) The ratio of pro-IL-1β/cleaved-IL-1β relative protein expression in neutralising antibody-treated samples to isotype control-treated samples. (D) The ratio of the caspase-1 activity in the supernatant (expressed as RLUs) from neutralising antibody-treated samples to isotype control-treated samples. (E) The levels of cytokine IL-1β in the culture supernatant were measured using ELISA. (F) The percentage of IL-1β inhibition in the supernatant in isotype control-treated and neutralising antibody-treated samples. (G) Isolated monocytes and autologous memory CD4+ T-cells from human PBMCs were co-cultured in the presence of anti-CD3 antibodies and the combination of neutralising antibodies (TNF-α Ab+CD40L Ab, TNF-α Ab+IFN-γ Ab, CD40L Ab+IFN-γ Ab) for 16 hours. The levels of cytokine IL-1β in the culture supernatant were measured using ELISA. (H) The percentage of IL-1β inhibition in single antibody-treated and combination antibody-treated samples. Data are represented as mean and SEM of independent donors in independent experiments. The P-value was assessed by students t-test or one-way ANOVA (* P≤ 0.05; ** P≤ 0.01; *** P≤ 0.001).

Our findings in immunoblotting analyses were validated further using ELISA to quantify IL-1β levels in the supernatant (Fig. 3E and F). To assess potential redundancy between TNF-α, CD40L, and IFN-γ in driving IL-1β secretion, we combined blocking antibodies. Dual inhibition of any two factors resulted in a significant reduction in IL1β secretion (Fig. 3G). Notably, dual blockade generally demonstrated greater efficacy in reducing IL-1β levels compared to single-factor inhibition (Fig. 3H). Although not statistically significant, a stronger inhibitory trend was observed with combined CD40L/IFN-γ blockade compared to CD40L inhibition alone. In summary, our data suggest that T cell-instructed IL-1β secretion by primary human monocytes is dependent on TNF-α, CD40L, and IFN-γ, but not FasL.

### T-cell-induced IL-1β secretion by human macrophages and dendritic cells

In murine models, T-cell-instructed IL-1β secretion has been observed in macrophages and dendritic cells (Jain et al., 2020). We investigated whether the mechanisms observed in primary human monocytes also apply to human macrophages and dendritic cells. Using GM-CSF-induced monocyte-derived macrophages (MDMs), M-CSF-induced MDMs, and IL-4 and GM-CSF-induced monocyte-derived dendritic cells (MDDCs), we first confirmed the induction of IL-1β secretion in these cells following stimulation with autologous memory CD4+ T cells (mCD4) (Fig. 4A). Next, we found that IL-1β production in T-cell-activated macrophages and dendritic cells depended on caspase-1 and required TNF-α, CD40L, and IFN-γ, but not FasL (Fig. 4B and C). These findings suggest that T-cell-induced IL-1β production is governed by shared mechanisms across different subsets of human myeloid cells.

**Figure 4.**
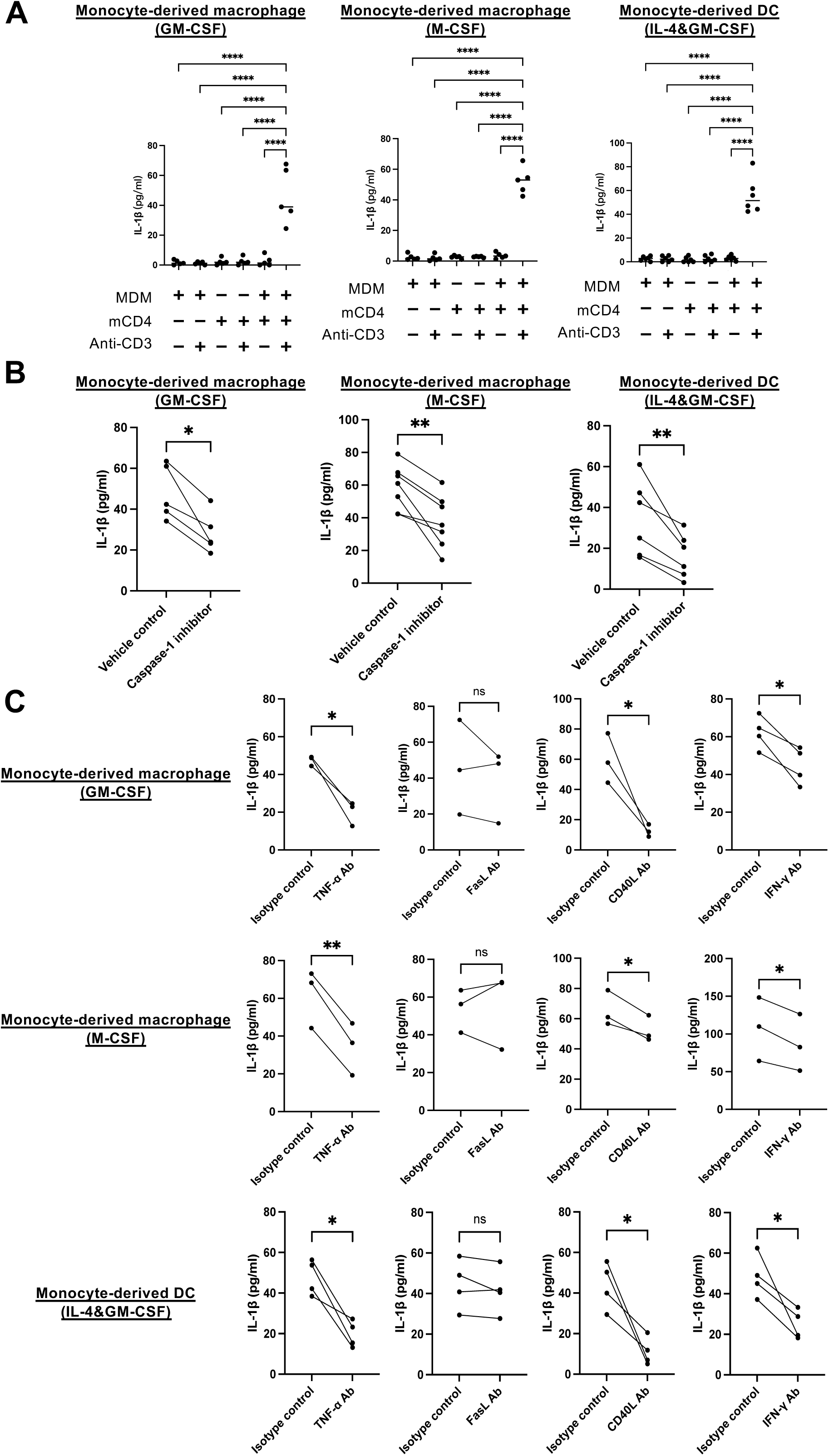
T-cell-mediated IL-1β secretion in human macrophages and dendritic cells is driven through TNF-α/CD40L/IFN-γ signaling and caspase-1 activation. (A) GM-CSF-induced MDM/M-CSF-induced MDM/MDDC and autologous memory CD4+ T-cells from human PBMCs were cultured alone or co-cultured in the absence/presence of anti-CD3 antibodies for 16 hours. The levels of cytokine IL-1β in the culture supernatant were measured using ELISA. (B) GM-CSF-induced MDM/M-CSF-induced MDM/MDDC were pre-treated with the caspase-1 inhibitor for 4 hours, followed by co-culture with autologous memory CD4+ T-cells and anti-CD3 antibodies for 16 hours. The level of IL-1β in the culture supernatant was measured by ELISA. (C) GM-CSF-induced MDM/M-CSF-induced MDM/MDDC and autologous memory CD4+ T-cells were co-cultured in the presence of anti-CD3 antibodies and neutralising antibodies against TNF-α, FasL, CD40L, and IFN-γ for 16 hours. The levels of cytokine IL-1β in the culture supernatant were measured using ELISA. Data are represented as mean and SEM of independent donors in independent experiments. The P-value was assessed by students t-test or one-way ANOVA (* P≤ 0.05; ** P≤ 0.01; *** P≤ 0.001).

### Concluding remarks

In this study, we demonstrate that activated T cells can induce IL-1β secretion by human monocytes, macrophages, and dendritic cells. This response involves the upregulation of pro-IL-1β mediated by TNF-α, CD40L, and IFN-γ, along with its cleavage by caspase-1. Interestingly, a prior study using murine cells also identified T-cell-instructed IL-1β production by myeloid cells but highlighted distinct mechanisms (Jain et al., 2020). Specifically, murine cells primarily relied on TNF-α (not CD40L) for pro-IL-1β upregulation, and caspase-8, rather than caspase-1, for IL-1β activation. These findings underscore that while T-cell-driven IL-1β responses are conserved across species, the underlying molecular pathways differ between humans and mice.

IL-1β production by myeloid cells is typically triggered by pathogen-associated molecular patterns (PAMPs) or damage-associated molecular patterns (DAMPs) and further amplified by danger signals such as ATP or reactive oxygen species (ROS). In this study, we demonstrate that activated T cells can drive both the upregulation of pro-IL-1β and its cleavage, highlighting a mechanism potentially relevant to the pathology of autoimmune diseases like rheumatoid arthritis (RA).

TNF-α blocking antibodies are a first-line treatment for several autoimmune diseases but benefit only about 50% of patients (Braun et al., 2015, Kumar et al., 2024, Lipsky et al., 2000). Our data reveal that the reduction of IL-1β achieved by TNF-α blockade is enhanced further through the co-inhibition of CD40L or IFN-γ. This suggests that bi- or tri-specific blocking antibodies targeting these pathways could potentially benefit patients who do not respond to TNF-α inhibition alone. Future experimental medicine studies are necessary to evaluate this hypothesis and evaluate its therapeutic potential.

In summary, we demonstrate that activated T cells induce IL-1β production in human myeloid cells through the upregulation of pro-IL-1β by TNF-α, CD40L, and IFN-γ, followed by caspase-1-mediated cleavage. T-cell-driven IL-1β responses are conserved between humans and mice, but they utilize distinct molecular pathways.

## MATERIALS AND METHODS

### Human subjects and cell isolation

Fresh leukocyte reduction system (LRS) cones from healthy donors (n=56) supplied by the NHS Blood and Transplant service. Human PBMCs were isolated from LRS cones by density gradient centrifugation on Histopaque-1077 (Sigma-Aldrich). Primary human monocytes, memory CD8+ T cells and memory CD4+ T cells were isolated from PBMCs using CD14 MicroBeads™, memory CD8+ isolation kit and memory CD4+ isolation kit according to the manufacturer’s instructions (Miltenyi Biotec), respectively. Human PBMCs, CD14+ monocytes, memory CD8+ T cells or memory CD4+ T cells were cultured in RPMI 1640 medium (Sigma) supplemented with 10% fetal bovine serum (FBS) and 1% GlutaMAX (Gibco) at 37°C in a humidified atmosphere containing 5% carbon dioxide.

### Monocyte-derived macrophages and dendritic cells preparation

Isolated monocytes were plated at 6 × 10^6^ cells per 12 mL in 100x20mm TC-treated culture dishes (Corning) in RPMI 1640 medium (Sigma) supplemented with 10% FBS and 1% GlutaMAX (Gibco). Monocytes were subsequently differentiated into GM-CSF-induced MDM and M-CSF-induced MDM by the addition of recombinant human GM-CSF (40 ng/mL, PeproTech) or recombinant human M-CSF (40 ng/mL, BioLegend Inc) for 6 days, respectively. For DC generation, recombinant human GM-CSF (40 ng/mL, PeproTech) and IL-4 (31 ng/mL, PeproTech) were added at the initiation of plated monocyte culture. On day 3, half of the culture medium was replaced with the fresh medium containing the same concentration of GM-CSF, M-CSF and IL-4. On day 6, cells were harvested for further experiments.

### Co-culture and in vitro treatment of monocytes/macrophages/DCs and T cells

Monocytes or monocyte-derived dendritic were cultured in non-tissue-culture-treated round-bottom 96-well plates (Corning) with memory CD4+ T cells at a ratio of 1∶3. Monocyte-derived macrophages and memory CD4+ T cells were co-cultured at 1:3 ratios in tissue-culture-treated flat-bottom 96-well plates (Falcon).

Monocytes/macrophages/DCs were preincubated with caspase-1 inhibitor (Ac-YVAD-cmk, InvivoGen, 20uM), caspase-8 inhibitor (Z-IETD-FMK, R&D Systems, 20uM) or neutralising antibodies against TNF-α (R&D Systems, 5μg/mL), FasL (R&D Systems, 10μg/mL), CD40L (BioLegend, 5μg/mL), and IFN-γ (R&D Systems, 5μg/mL) before the co-culture. For stimulation assays, monocytes were stimulated with either LPS (Enzo, 100ng/ml) plus nigericin or purified anti-human CD3 antibody (clone OKT3, BioLegend) for 16 hours. Memory CD4+ T cells and memory CD4+ T cells were stimulated with purified anti-human CD3 antibody (clone OKT3, BioLegend) for 16 hours.

#### siRNA silencing in human primary monocytes

Primary human monocytes were transfected with 20nM of control non-targeting siRNA or siRNA targeting human CASP1 and CASP8 (Dharmacon) using HiPerFect reagent (QIAGEN) following the instructions provided by the manufacturer in 24-well plates. After 72 hours, the cells were collected for subsequent experiments.

#### Enzyme-linked immunosorbent assays (ELISA)

IL-1β in cell culture supernatants were assayed using a standard ELISA kit (Invitrogen) according to the manufacturer’s instruction. The Multiskan Ascent ELISA reader was employed to read the ELISA plates at a test wavelength of 450 nm and a background wavelength of 570 nm. Then, the standard curve of ELISA was plotted using Microsoft Excel software, while the corresponding concentrations of samples were calculated using the absorbance value and standard curve.

#### Intracellular cytokine staining (ICS)

After washing in PBS with 1% FBS, cells were surface stained for 30 minutes at 4°C with live/dead kit (Invitrogen), anti-CD3-Brilliant Violet 786 (Biolegend), and anti-CD14-Brilliant Violet 711 (Biolegend). Cells were then fixed and permeabilized in the Transcription Factor Buffer Set (562574, BD Biosciences) for intracellular cytokine staining. FACS anti-human antibodies for IL-1 beta (clone CRM56 eBioscience) were used. Stained cells were analysed using a BD LSR Fortessa (BD Biosciences) and FlowJo analytical software (Treestar).

#### Western blotting

Cells were lysed using RIPA buffer supplemented with protease inhibitors (Sigma). Protein concentrations were quantified using the bicinchoninic acid (BCA) assay. Total proteins were electrophoresed through the NuPAGE 4-12% Bis-Tris Protein Gels (Life Technologies), transferred to the nitrocellulose membranes (Thermo Fisher Scientific) and analysed by immunoblotting. The primary antibodies used were rabbit anti-IL-1β (1:1000 dilution, Cell Signalling, D3U3E), rabbit anti-cleaved-IL-1β (1:1000 dilution, Cell Signalling, D3A3Z), rabbit anti-caspase-1 (1:1000 dilution, Cell signalling, D7F10), mouse-anti-caspase-8 (1:1000 dilution, Cell signalling, 1C12) and mouse anti-β-Actin (1:1000 dilution; Sigma-Aldrich, AC-15). The secondary antibodies used were IRDye 800CW goat anti-mouse IgG and IRDye 680RD goat anti-rabbit IgG (1:10000 dilution, LI-COR Biosciences). The bands on the membranes were scanned using the LI-COR Odyssey Imager (LI-COR, USA) and analysed with Image Studio software (LI-COR Biosciences) and Image J software (Wayne Rasband, NIH, USA).

#### Caspase-1 activity assay

The activity of caspase-1 was measured using the Caspase-Glo 1 Inflammasome Assay (Promega) following the manufacturer’s instructions. Cell supernatants were collected and incubated with the reconstituted Caspase-Glo® reagent (Z-WEHD-aminoluciferin caspase-1 substrate) in 96-well plates at room temperature for 1 hour. The luminescence value was measured using a microplate reader (Fluostar Omega, BMG Labtech).

#### Statistics

All statistical analyses and summarized graphs were performed using GraphPad Prism version 9.4.1. Experiments were repeated at least three times with similar results. The level of statistical significance was assessed by the two-tailed Student’s t-test (between two group comparisons) or the ANOVA (between multiple group comparisons). The differences were considered statistically significant at P<0.05*, P<0.01**, P<0.001***.

#### Study approval

Venous blood and synovial fluid were obtained under protocols approved by the Oxford Research Ethics committee (Ethics reference number 06/Q1606/139).

#### Data availability

The data supporting the findings of this study are available within the article and the main figures or its supplementary materials. Raw western blot images demonstrated in the figure sand used to generate plots are available upon request.

## AUTHOR CONTRIBUTIONS

J.Z., Z.L. and L.C. designed the research studies, J.Z. and Z.L. were conducting experiments, J.Z. and Z.L. were acquiring data, J.Z. and L.C. were analysing data. J.Z., Z.L., B.M.K. and L.C. were all contributing to writing the manuscript.

## ACKNOWLEDGEMENTS

This work was funded by a Versus Arthritis career development award to LC 22053.

Z.L. and B.M.K. were supported by the Chinese Academy of Medical Sciences (CAMS) Innovation Fund for Medical Science (CIFMS), China (grant number: 2018-I2M-2-002), awarded to B.M.K.

## FIGURE LEDENDS

**Figure S1.**
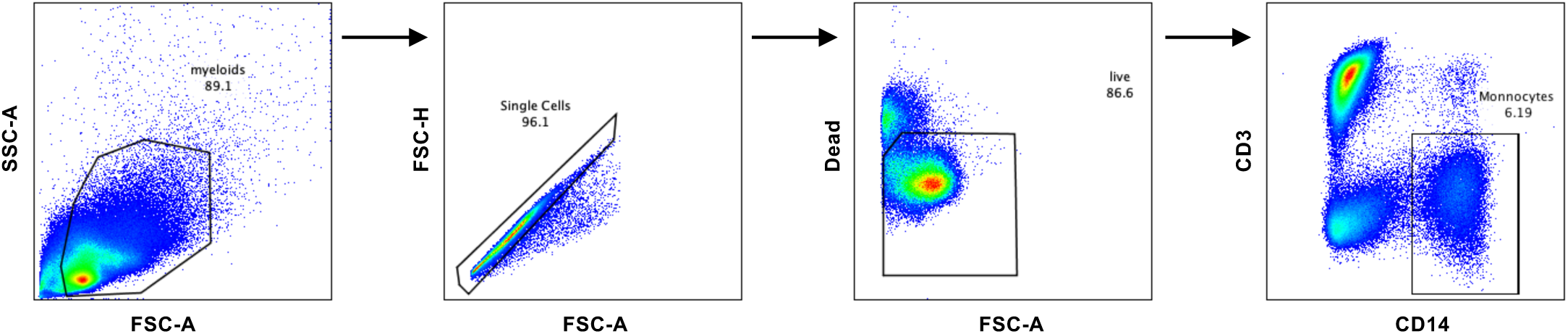
The gating strategy for Figure 1C to identify CD14+ monocytes. Isolated human PBMCs were stimulated with LPS (100ng/mL) or cultured with memory CD4+ T cells in the presence or absence of anti-CD3 antibody for 16 h. Myeloid cells were gated on singlet cells followed by live cells. Monocytes were defined as CD3−CD14+ cells.

**Figure S2.**
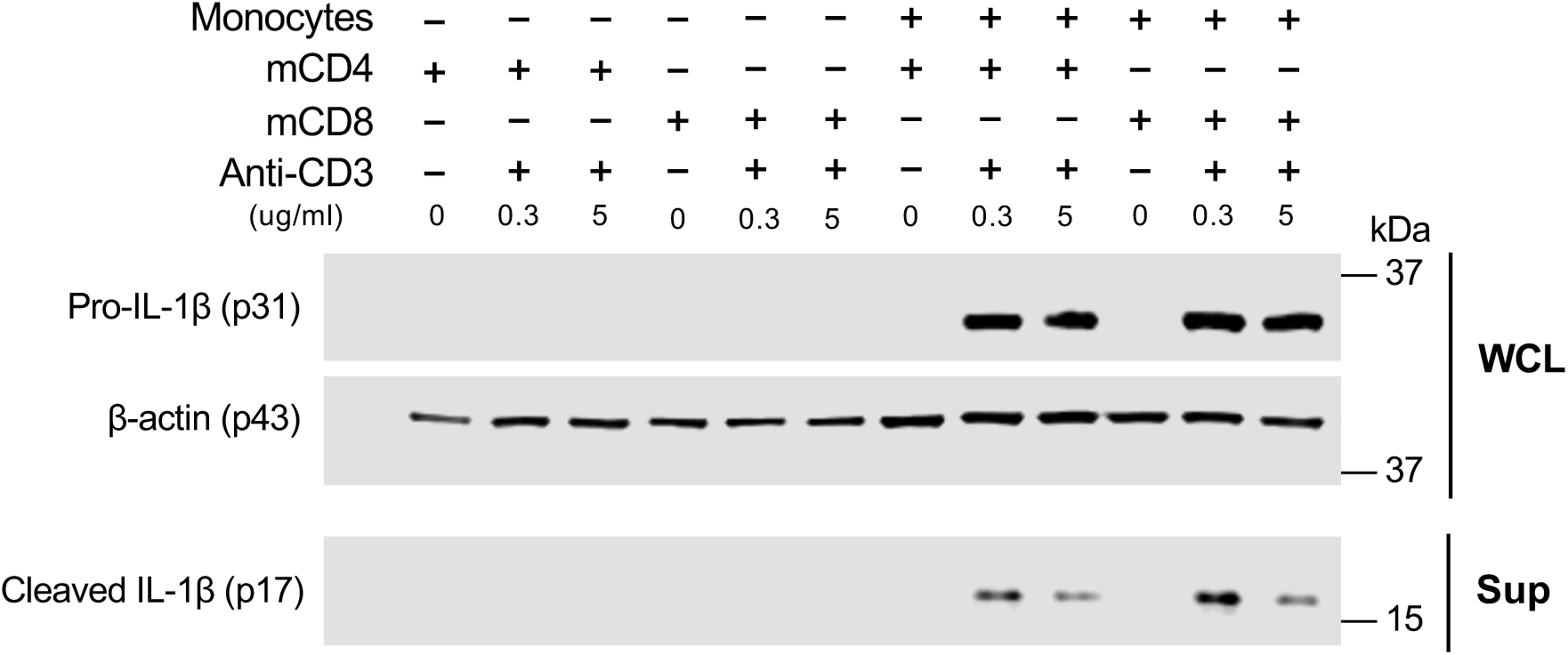
Both activated CD4+ and CD8+ T cells trigger the production and secretion of IL-1β in monocytes. Isolated monocytes, autologous memory CD4+ T-cells or memory CD8+ T-cells from human PBMCs were cultured alone or co-cultured in the absence/presence of anti-CD3 antibodies for 16 hours. Immunoblotting analysis of pro-IL-1β, β-actin, and cleaved-IL-1β in the whole cell lysates or the supernatants.

**Figure S3.**
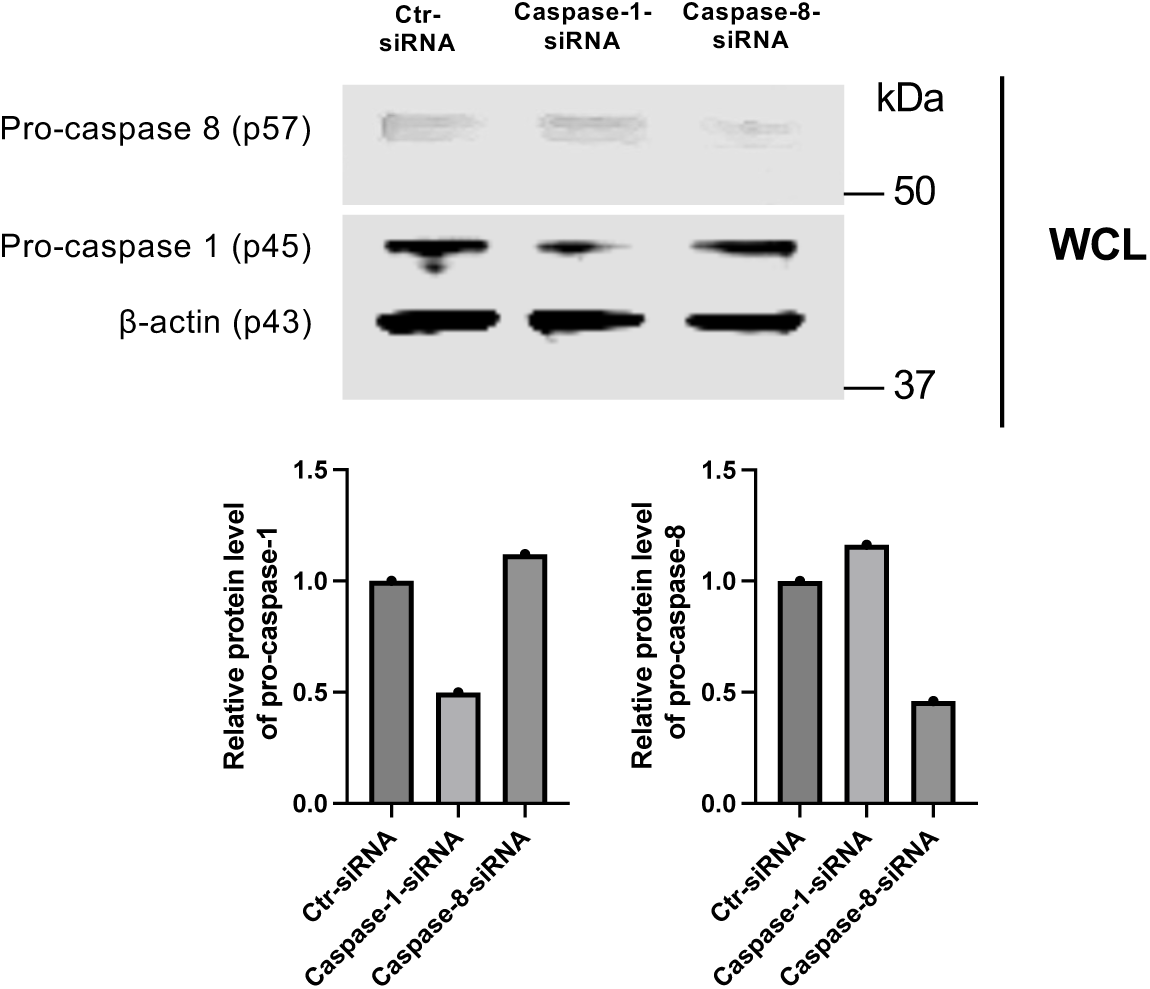
Efficiency of siRNA-mediated knockdown for Caspase-1 and Caspase-8. Isolated monocytes from human PBMCs were transfected with non-targeting control siRNA, CASP1 siRNA, or CASP8 siRNA before co-culture with autologous memory CD4+ T cells and anti-CD3 antibodies for 16 hours. Immunoblotting analysis of cell lysate from transfected monocytes with anti-caspase-1, anti-caspase-8, and anti-β-actin antibodies. The bar graphs present the relative protein levels of caspase-1 and caspase-8 in transfected monocytes (GraphPad).

